# Cortical cell size regulates root metabolic cost

**DOI:** 10.1101/2023.08.18.553921

**Authors:** Jagdeep Singh Sidhu, Jonathan P. Lynch

## Abstract

It has been hypothesized that vacuolar occupancy in mature root cortical parenchyma cells regulates root metabolic cost and thereby plant fitness under conditions of drought, suboptimal nutrient availability, and soil mechanical impedance. However, the mechanistic role of vacuoles in reducing root metabolic cost was unproven. Here we provide evidence to support this hypothesis. We first show that root cortical cell size is determined by both cortical cell diameter (CCD) and cell length (CCL). Significant genotypic variation for both CCD (∼1.1 to 1.5- fold) and CCL (∼ 1.3 to 7-fold) was observed in maize and wheat. GWAS and QTL analyses indicate CCD and CCL are heritable and under independent genetic control. We identify candidate genes for both phenes. Empirical results from isophenic lines contrasting for CCD and CCL show that increased cell size, due to either CCD or CCL, is associated with reduced root respiration, root nitrogen content, and root phosphorus content. *RootSlice*, a functional-structural model of root anatomy, predicts that an increased ratio of vacuolar to cytoplasmic volume causes reduced root respiration and tissue nutrient content. Ultrastructural imaging of cortical parenchyma cells with varying CCD and CCL confirms the *in-silico* predictions and shows that an increase in cell size is correlated with increased vacuolar volume and reduced cytoplasmic volume. Phylogenetic analysis of terrestrial plants reveals that CCD has not significantly changed throughout plant evolution. Vacuolar occupancy and its relationship with CCD/CCL merits further investigation as a phene for improving crop adaptation to edaphic stress.

**Significance Statement:** Cortical cell size is an important phene determining root metabolic cost, but the underlying physiological mechanism is unclear. Here, using *in silico* and empirical approaches, we provide evidence that supports the hypothesis that vacuolar occupancy in cortical parenchyma cells regulates root metabolic cost. We also show that vacuolar occupancy is associated with cortical cell diameter and cell length, phenes that are under distinct genetic control and hold the potential for improving crop yields under edaphic stress.

## Introduction

Drought stress and low soil fertility are primary limitations to global crop production (1, 2). The severity of these constraints is projected to worsen with ongoing climate change and soil degradation (3–6). To achieve more resilient agricultural systems, we must develop novel crop varieties that can acquire and use limited resources more efficiently (7, 8). Understanding resource partitioning, especially under edaphic stress conditions such as drought, low nitrogen, and low phosphorus conditions, is crucial to identify varieties with improved fitness. Under nutrient or water stress conditions, the root system can consume up to 50% of daily photosynthates for the maintenance of living cells (9, 10). Plants also invest a significant portion (>25%) of scarce nutrients like nitrogen (N) and phosphorus (P) for the maintenance of root tissue (11–13). Therefore, phenes (‘phene’ is the fundamental unit of the phenotype (14–16)) that reduce the nutrient and carbon demand of the root system provide an opportunity for efficient and extensive soil exploration (17–19).

Root anatomy is an important determinant of root metabolic cost (19, 20). Over the past decade, several root anatomical phenes regulating root metabolic cost and thereby improving plant fitness under edaphic stress have been discovered (20). For example, increased root cortical aerenchyma (RCA) in maize and common bean, reduced cell file number (CCFN) in maize, increased root cortical senescence (RCS) in barley, reduced root secondary growth (RCS) in common bean, and increased cortical cell size (CCS) in maize and wheat, independently, lead to reduced root metabolic cost and improve plant performance under edaphic stress (21–26). Among the known phenes regulating root metabolic cost, CCS is one of the most important and fundamental yet is relatively underexplored and mechanistically less understood. Larger root CCS was associated with reduced root metabolic cost by three independent studies on maize and wheat (23, 24). Larger CCS in maize reduced root respiration by 59%, increased rooting depth by 25%, and increased yield by ∼2-fold under drought stress as compared to smaller CCS (23). Increased CCS improved maize growth under suboptimal nitrogen availability *in silico* (27). Larger CCS in wheat grown in mechanically impeded soils reduced root respiration by ∼50% (24). Larger CCS in maize is also associated with improved penetration of hard soils (28). These studies show that large CCS reduces root metabolic cost, enables deeper soil exploration, and increases water and nutrient acquisition, hence improving plant fitness and yield under limited resources. However, the mechanism by which larger CCS reduces root metabolic cost is unclear.

The influence of cell dimensions on physiological functions is not unique to plants and has been the focus of research in several biological fields. The relationship between cell size and metabolic cost has been explored in detail in animal species. For example, increased cell size correlates with reduced metabolic rate (corrected for body mass) in amphibians (29), eyelid geckos (30), carabid beetles (31), birds (32), and mammals (33). The underlying cause of the negative relationship between cell size and metabolic rate is still debatable, in fact, a positive relationship between cell size and the metabolic rate has been observed in some animal tissues (34, 35). Glazier (2022) highlights several plausible causes for a negative correlation between cell size and metabolic rate in animal tissues (36). This includes the involvement of reduced resource supply and/or demand in larger cells, lower metabolic costs of ionic regulation, slower cell multiplication and somatic growth, and larger intracellular deposits of metabolically inert materials in some tissues (36). Most of these explanations do not translate to plant cells given the obvious intracellular differences between animal and plant cells. However, intracellular deposition of metabolically inert material seems to be a common phenomenon. For example, in animal cells, metabolic efficiency can be achieved by the deposition of fat in animal adipose tissue cells (which are significantly larger than other cell types) (37, 38). In plants, the deposition of low-energy materials such as lignin, cellulose, starch, and water can similarly reduce tissue metabolic rate (39, 40).

Plant cells deploy vacuolar expansion to enlarge in an energy-efficient manner (41–43). Depending on cell type and ontogeny, more than 90% of plant cell volume can be occupied by vacuolar compartments. The space-filling role of vacuoles and their reduced metabolic cost compared to the cytosol allows rapid plant cell expansion (44). This energy-efficient cell expansion mechanism may be a factor in the large size of plant cells, which are on average ∼ 80 to 350 times larger than high-metabolic rate endothermic birds and mammals, whereas medium-metabolic rate ectothermic vertebrates have cells with intermediate volumes (36).

It has been proposed that the size of root cortical parenchyma cells driven by vacuolar occupancy can regulate the metabolic cost of soil exploration and hence the capture of water and mineral nutrients (45). The hypothesis states that roots with larger CCS are energetically cheap due to an increase in the ratio of vacuolar to cytoplasmic volume in root tissue. The negative relationship between root CCS and root metabolic cost was corroborated in maize and wheat (23, 24), as discussed earlier. However, the involvement of vacuoles was not explicitly shown by either of these studies. Indirect evidence supporting the importance of vacuoles in reducing root metabolic cost can be gathered from studies comparing different root development stages (42). Root meristematic cells are comparatively small but have higher metabolic activity compared to mature root cortical cells which have large central vacuoles (42, 46). Similarly, xylem parenchyma and phloem companion cells tend to have reduced vacuolar volume and increased cytosolic content to support the energy-intensive process of transporting ions and solutes (47, 48). Besides the study involving the comparison of different tissue types, which are confounded by other factors, no direct evidence has been provided to show how vacuole occupancy can regulate root metabolic costs. The role of vacuoles in regulating root metabolic cost can be explicitly shown by comparing genotypes with contrasting root CCS.

Root CCS is determined by both cortical cell length (CCL) and cortical cell diameter (CCD) (23, 49). Both CCD and CCL are important for soil exploration. CCL determines the root elongation rate and its interplay with cell division determines root length (50). CCD determines the cross-sectional area of a cell and its interaction with root cortical cell file number and stele diameter regulate root diameter (51). Both CCD and CCL, however, are associated with vacuolar occupancy, thus providing a system to decipher the role of CCS and vacuoles in controlling root metabolic costs.

In this study, we primarily use maize and wheat, primary global crops, as models to test the hypothesis that increased vacuole:cytoplasm ratio per unit cortical volume is the underlying cause of the association of CCS and root metabolic cost. Our approaches include *in silico* modeling, analysis of genotypes naturally contrasting for CCD and CCL, and electron microscopy. We first show that plants can use vacuolar expansion to alter cell size either by changing cortical cell diameter (CCD) or cortical cell length (CCL). Additionally, we explore the underlying genetics and evolutionary changes in CCS.

## Results

### Natural variation for CCD and CCL

The maize IBM RIL (Recombinant Inbred Line) population, maize WiDiv diversity panel, and wheat Watkin collection were utilized to assess natural genotypic variation for CCD and CCL. In all three populations, CCD approximately follows a Gaussian distribution with a slight skew to the right in the WiDiv panel and wheat Watkin collection (Figure 1A). In the IBM RIL population, we observed 1.5-fold variation (SD = 7.7 µm) in CCD, ranging from 25.1 to 63.9 µm, and centered at a median CCD of 40.3 µm (Figure 1A). In the WiDiv diversity panel, we see a 1.6- fold variation (SD = 4.6 µm) in CCD, ranging from 19.4 µm to 50.2 µm and a median of 29.2 µm (Figure 1A). In the wheat Watkin collection, we find 1.2-fold variation (SD = 3.2 µm) in CCD, with a median CCD of 24.5 µm ranging from 15.3 µm to 33.1 µm (Figure 1A).

**Figure 1.**
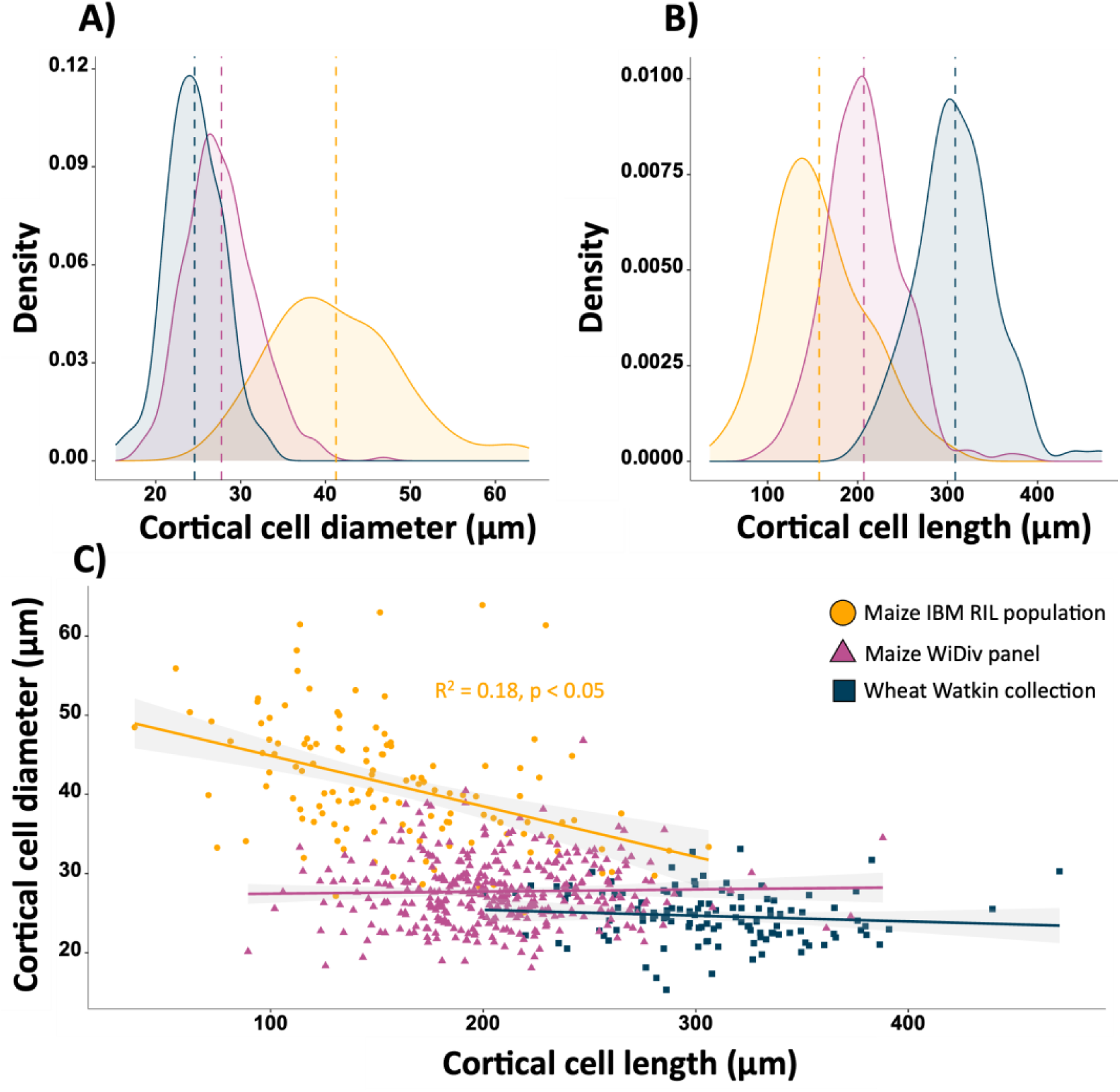
Natural variation in cortical cell diameter (CCD) and cortical cell length (CCL) in maize and wheat. Density plot showing variation for CCD (A) and CCL (B) in the maize WiDiv population, maize IBM population, and wheat Watkin collection. Relationship between CCD and CCL in all three panels, R^2^ and p-value only reported for the significant (p ≤ 0.05) relationships.

We observe similar trends for CCL. In all three populations, CCL follows a Gaussian distribution with slight right skewness in the WiDiv and IBM population (Figure 1B). In the IBM RIL population, the median CCL is 153.2 µm ranging from 36 to 306 µm (7.5-fold variation, SD = 51.2 µm) (Figure 1B). The median CCL in the WiDiv panel is 206.1 µm varying from 89.4 µm to 388 µm (3.4-fold variation, SD = 43.1 µm) (Figure 1B). In the wheat Watkin collection, we observe 1.3-fold variation (SD = 44.6 µm) in CCL, ranging from 200.7 to 471.1 µm and centered at a median CCL of 305.6 µm (Figure 1A).

Within the WiDiv panel and wheat Watkin collection, we observed no significant relationship between CCD and CCL (Figure 1C), however, a significant negative correlation (R^2^ = 0.18, p ≤ 0.05) between CCD and CCL was observed within the IBM population. Despite the negative correlation between CCL and CCD within the IBM population, we were able to identify isophenic genotypes contrasting solely for CCD and CCL (Figure 2). IBM-365 is a smaller CCD genotype with an average CCD of 26.2 µm while IBM-30 is a larger CCD genotype with an average CCD of 34.0 µm (Figure 2A and B). IBM-111 is a longer CCL genotype with an average CCL of 205 µm while IBM-177 is a shorter CCL genotype with an average CCL of 158.7 µm (Figure 2C and 2D).

**Figure 2.**
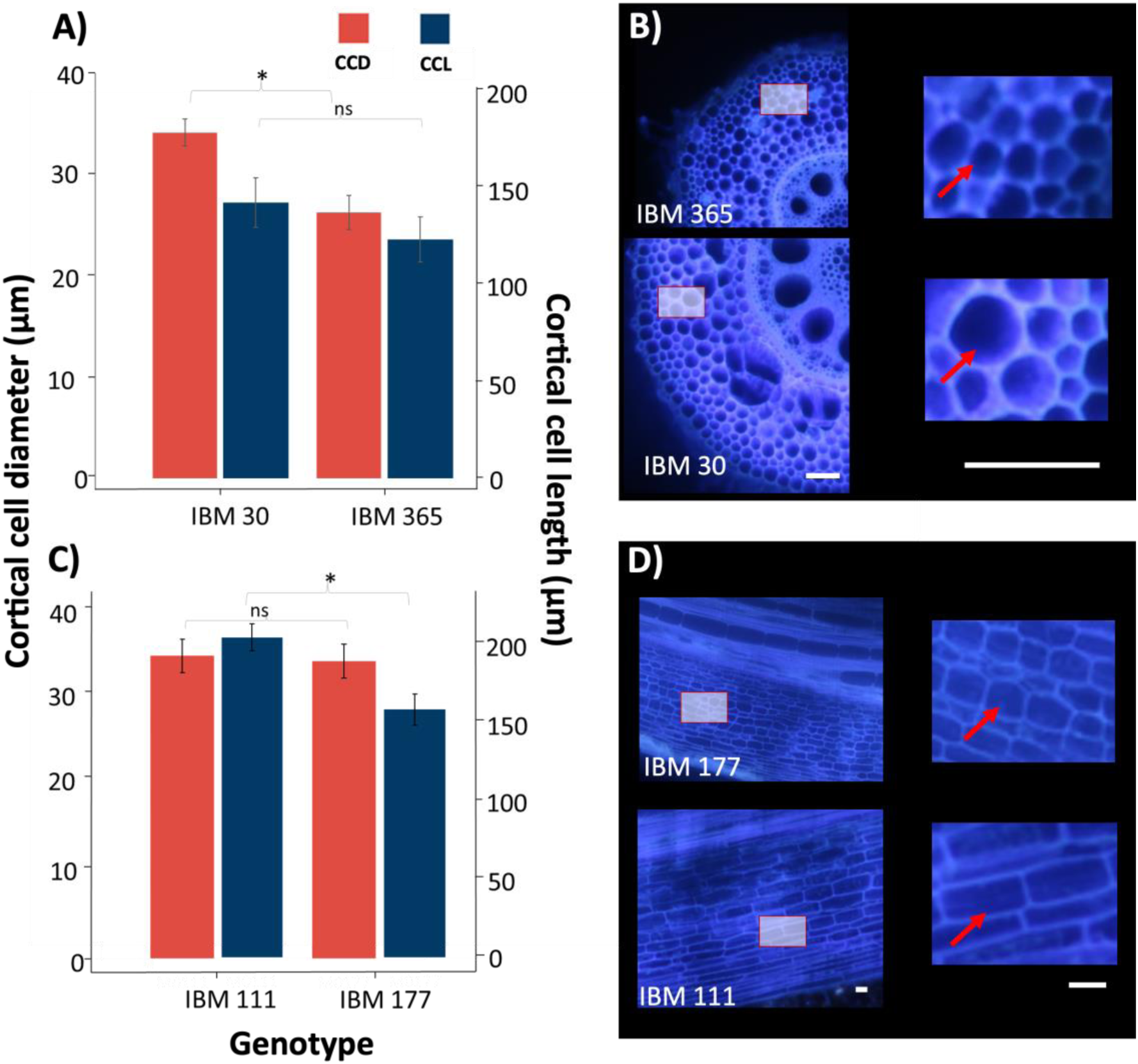
Contrasting genotypes for cortical cell diameter (CCD) and cortical cell length (CCL). IBM 30 and IBM 365 significantly differ for CCD (A), IBM 111 and IBM 177 significantly differ for CCL only (C). Root cross sections of fourth whorl roots of genotypes contrasting for CCD (B) and longitudinal sections second whorl roots of genotypes contrasting for CCL (D). Red arrows in each inset points to a representative cell for the respective genotype. Scales in C and D = 100 µm. Images in panel B and D were acquired using laser ablation tomography.

### Empirical data shows increased CCD and CCL is associated with reduced root respiration, nitrogen and phosphorus content

An increase in cell size, either longitudinally or cross-sectionally, is associated with decreased root respiration and nutrient content (Figure 3). The mature fourth whorl root segment of IBM-30, a genotype with larger CCD, respires at an average respiration rate of 7 nmol CO_2_ g^-1^ sec^-1^, which is 50 % less than the average respiration rate of IBM-365, a genotype with smaller CCD (Figure 3A). The average root tissue nitrogen concentration of a larger CCD genotype is 1.055 %, which is 20 % less than a genotype with smaller CCD (Figure 3B). Similarly, the average root tissue phosphorus content of a genotype with large CCD is 0.24 %, which is 42% less than a genotype with smaller CCD (Figure 3C).

**Figure 3.**
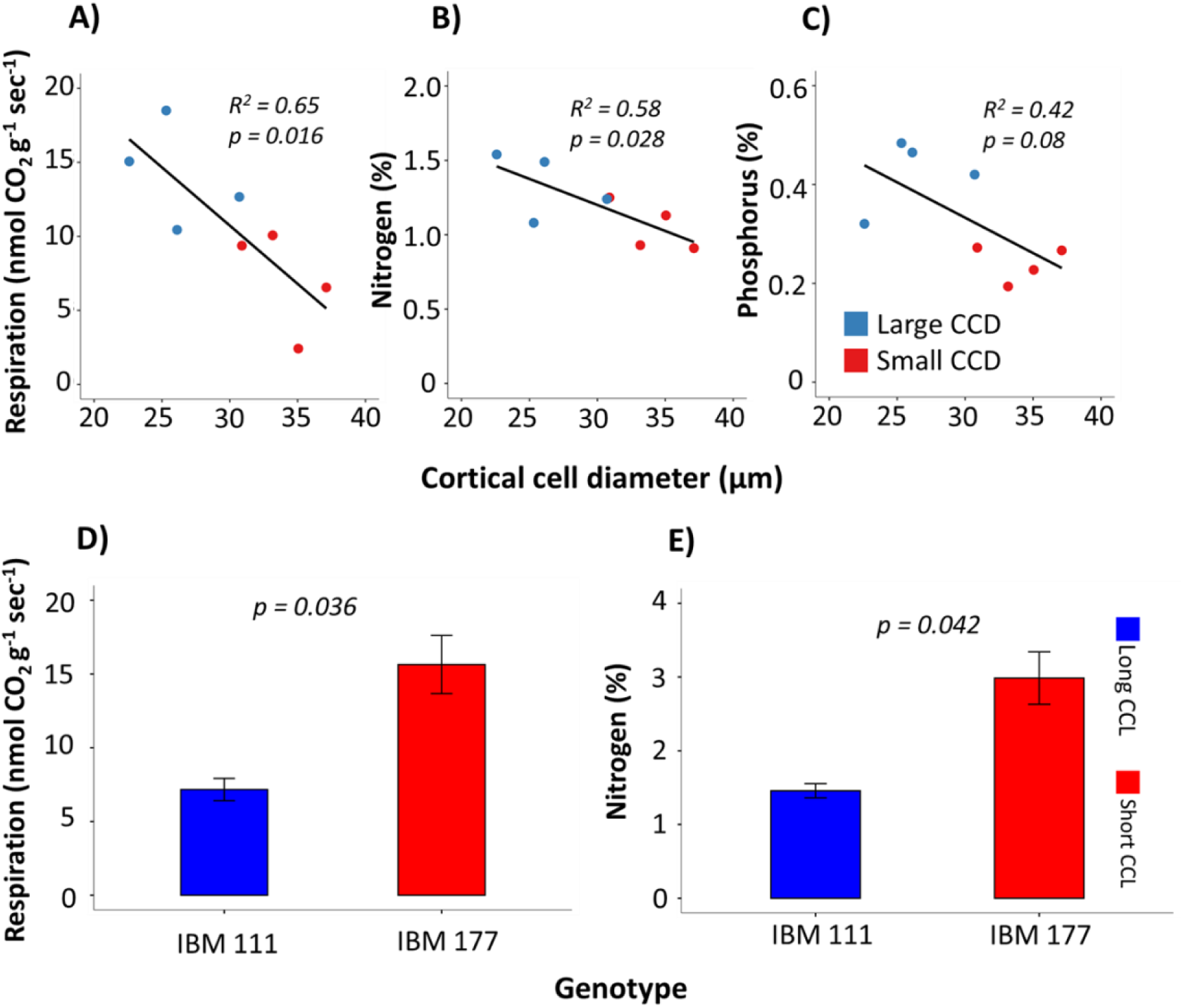
Cortical cell size influences root respirations and root nutrient content. Increase in cortical cell diameter (CCD) is associated with reduction in root respiration (A), nitrogen content (B) and phosphorus content (C). Genotype with longer cortical cells have lower root respiration (D) and root nitrogen content (E) compared to shorter cortical cell length genotype. n = 8 for A, B, and C; n = 6 for D; and n = 7 for E.

The average root respiration of a shorter CCL genotype (IBM-177) is 15 nmol CO_2_ g^-1^ sec^-1^, whereas a longer CCL genotype (IBM-111) respires at 7 nmol CO_2_ g^-1^ sec^-^1 (Figure 3D). Thus, IBM-111 respires at a 53% lower rate compared to IBM-177. In parallel, the average root nitrogen of IBM-111 is 1.45 % which is 41 % less than the root nitrogen content of IBM-177 (Figure (3E).

### *RootSlice* predicts vacuolar occupancy in larger cells is linked to reduced root respiration and nutrient content

For a simulated maize nodal root segment (Figure 4A and B), increasing cortical cell diameter from 10 to 25 µm reduces mitochondrial density by 51 % (Figure 1.5E), respiration by 49 % (Figure 4F), nitrogen content by 35 % (Figure 4G), and phosphorus content by 27 % (Figure 4H). *RootSlice* shows that this reduction in respiration and nutrient content is due to decreased cytoplasmic volume (by 51 %, Figure 4C) and a corresponding increase in vacuolar volume (by 17 %, Figure 4D) per unit cortical volume.

**Figure 4.**
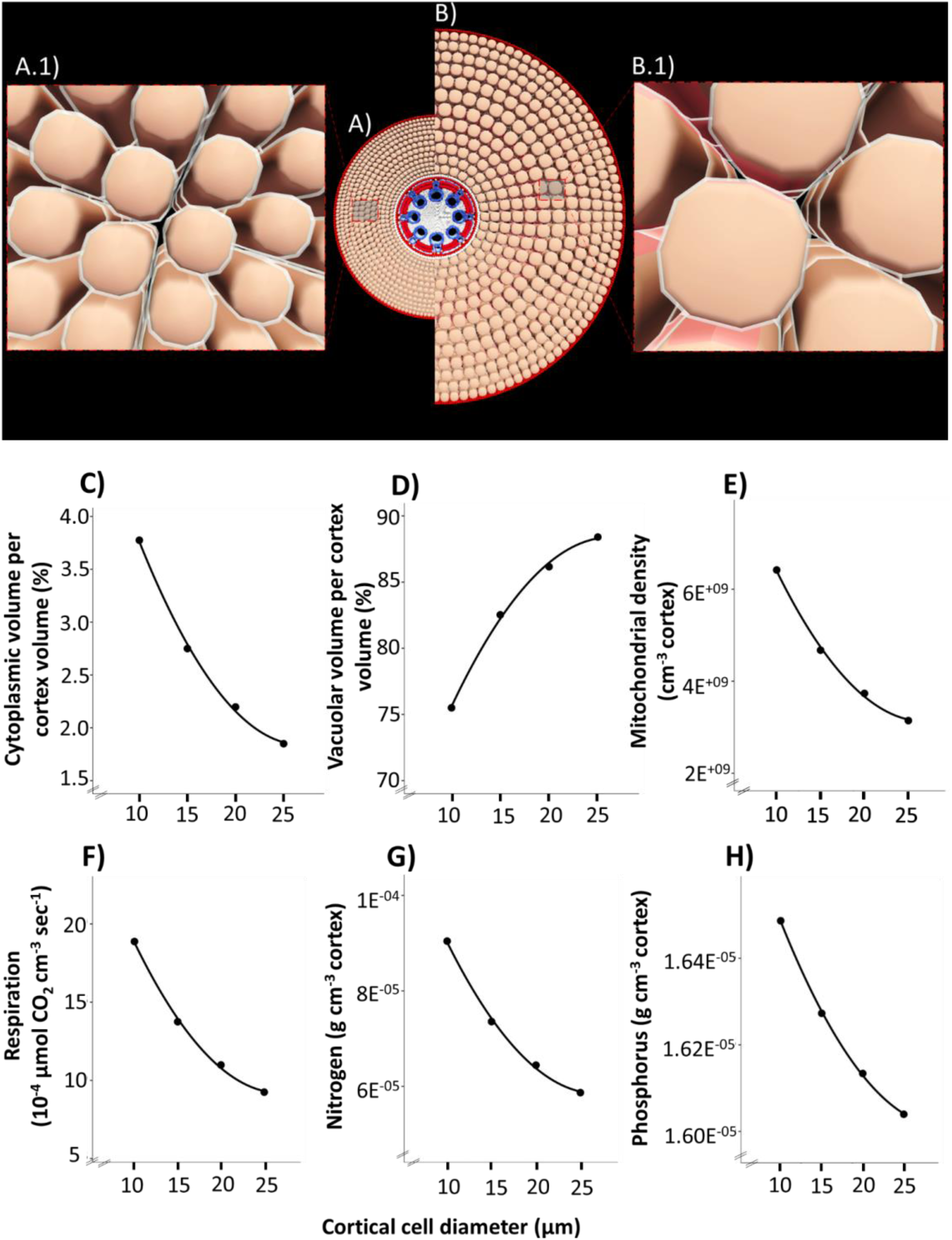
*RootSlice* results showing the effect of cortical cell diameter on root resource content. A) Simulated cross section with an average 10 µm cortical cell diameter and A.1 as an inset, B) average cortical cell diameter of 25 µm with B.1 as an inset. Color codes in the insets: cell wall (grey), plasma membrane (green), and vacuole (red). Effect of cell diameter on A) cytoplasmic volume, B) vacuolar volume, C) mitochondrial density, D) respiration, E) nitrogen, and F) phosphorus.

**Figure 5.**
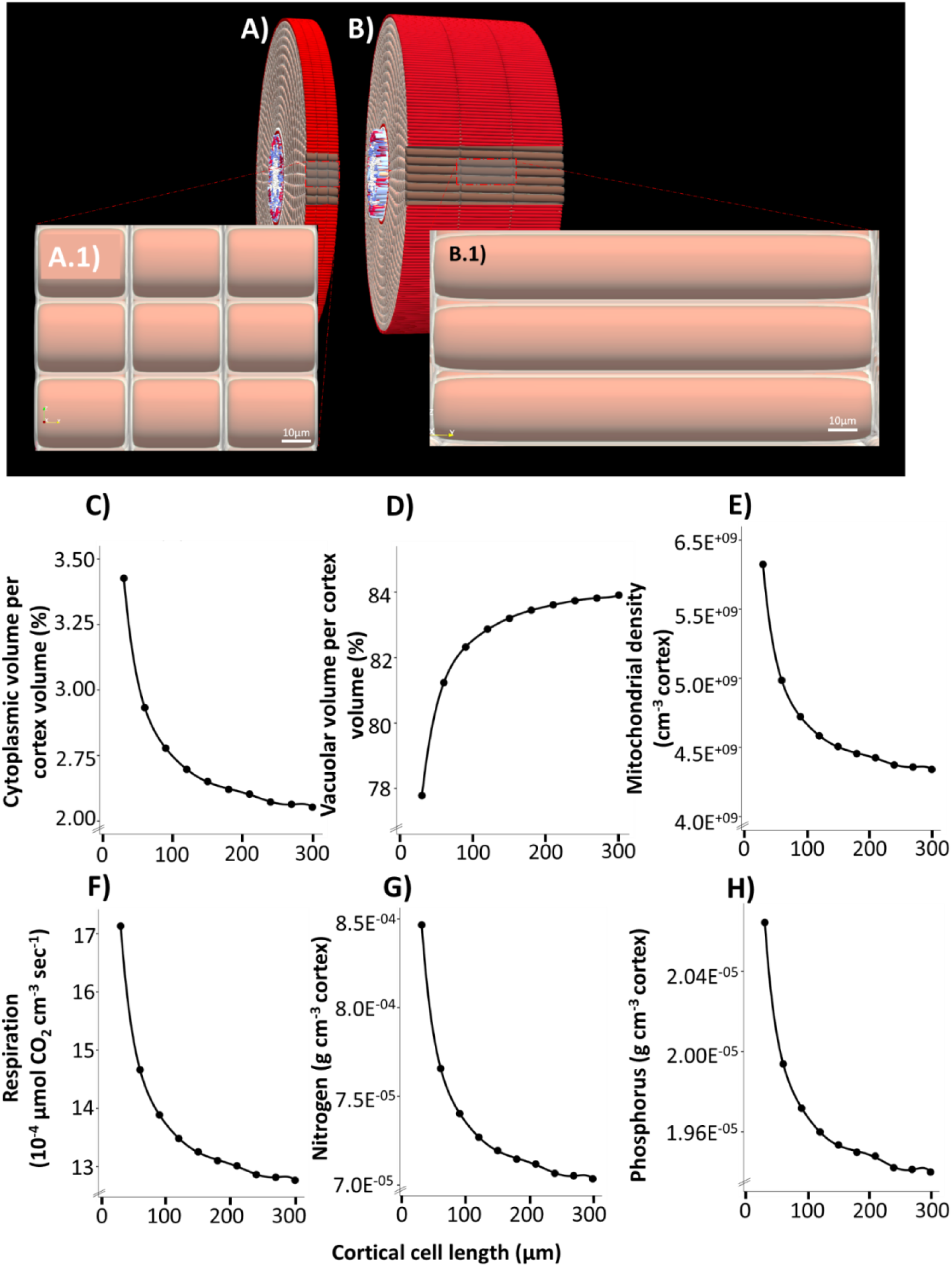
*RootSlice* simulations of different cortical cell lengths. A) Simulated cross section with an average 30 µm cortical cell length and A.1 as an inset, B) average cortical cell length of 150 µm with B.1 as an inset. Color codes in the insets: cell wall (grey), plasma membrane (green), and vacuole (red). Figure 8. *RootSlice* results showing the effect of cortical cell length on C) cytoplasmic volume, D) vacuolar volume, E) mitochondrial density, F) respiration, G) nitrogen, and H) phosphorus.

Greater CCL also reduces root metabolic cost *in silico*. Increased cell length (Figure 5A and B) from 30 to 300 µm reduces mitochondrial density by 14% (Figure 5E), respiration by 14% (Figure 5F), nitrogen content by 9% (Figure 5G), and phosphorus content by 5% (Figure 5H). *RootSlice* again shows that a reduction in cytoplasmic volume (by 14 %, Figure 5C) and an increase in vacuolar volume (by 4 %, Figure 5D) is the underlying cause for reduced respiration and root nutrient content.

### Greater CCD and CCL associated with greater vacuolar occupancy

Among other imaging techniques including TEM, two-photon microscopy SBFSEM provided the most detailed and well-preserved ultrastructural imaging. SBFSEM shows that independent of cell size, the average distance between the plasma membrane and tonoplast is approximately 0.2 µm. Detailed imaging of mature root cortical cells of different genotypes contrasting for CCD and CCL reveals that more than 90% of the cell volume is occupied by vacuolar compartments (Figure 6). An increase in cell size, shown here as cross-sectional surface area, is closely tracked by vacuolar occupancy and only a small increase in cytoplasmic area. With every 1 µm^2^ increase in cell cross-sectional area, there is a 0.86 µm^2^ increase in vacuolar area and only 0.14 µm^2^ increase in cytoplasmic area.

**Figure 6.**
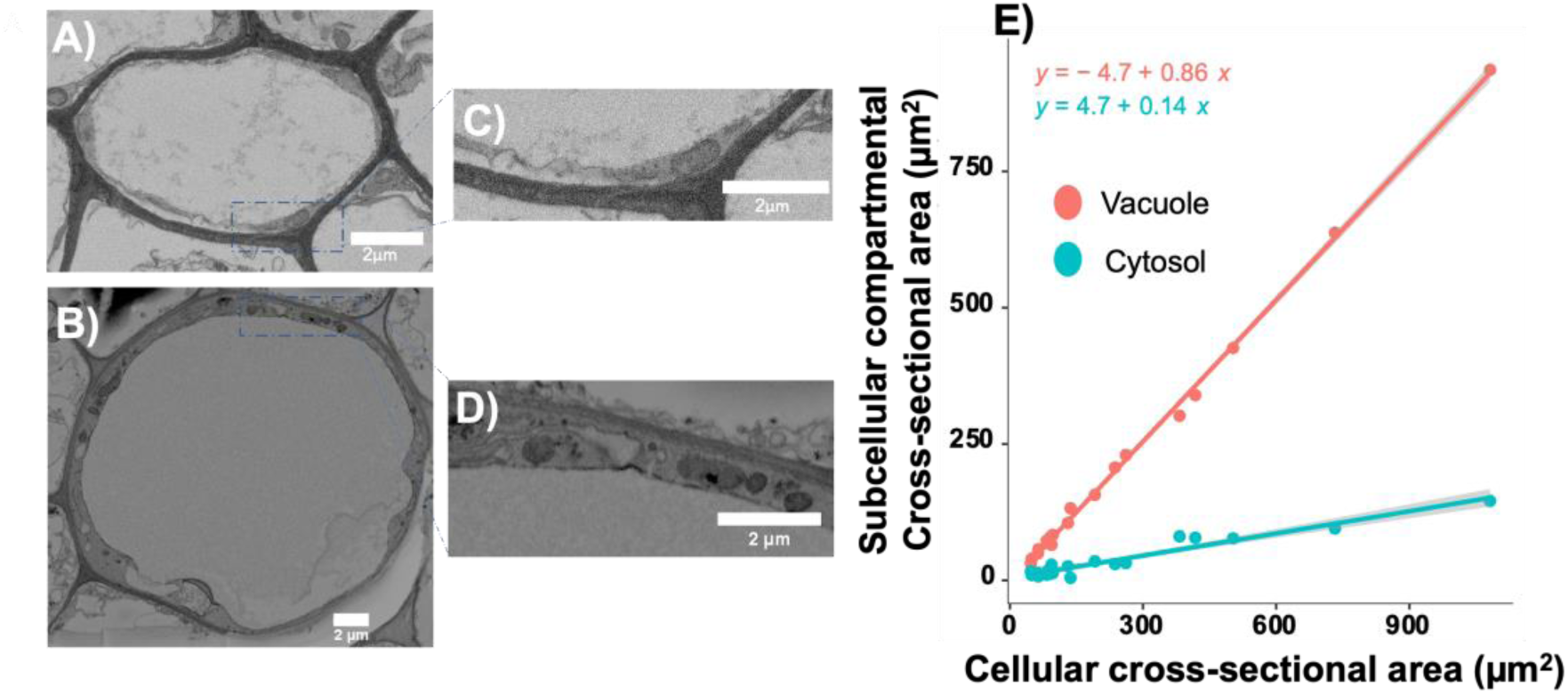
Serial block facing scanning electron microscopy (SBFSEM) imaging showing increased cell size is linked with increased vacuolar occupancy. Scan of cortical cell of IBM 365 (A, C), and IBM30 (B, D). Relationship between cell surface area and subcellular compartment surface area including vacuolar and cytosolic compartments.

### Genetic analysis of CCD and CCL in maize

QTL analysis of the IBM (intermated B73XMo17) population reveals a significant QTL (named *QTL_CCL_7*) linked to CCL on chromosome number 7 (Figure 7A). SNP m770 and m794 demarcate the *QTL_CCL_7* with m788 being the most significant SNP. Genotypes with homozygous BB allele of m788 have an average of 170 µm CCL, whereas genotypes with homozygous AA allele have an average CCL of 140 µm, and the heterozygous “AB” pool has the shortest cell length with 120 µm. *QTL_CCL_7* explains 11.05 % of the variation observed for CCL in the IBM mapping population.

**Figure 7.**
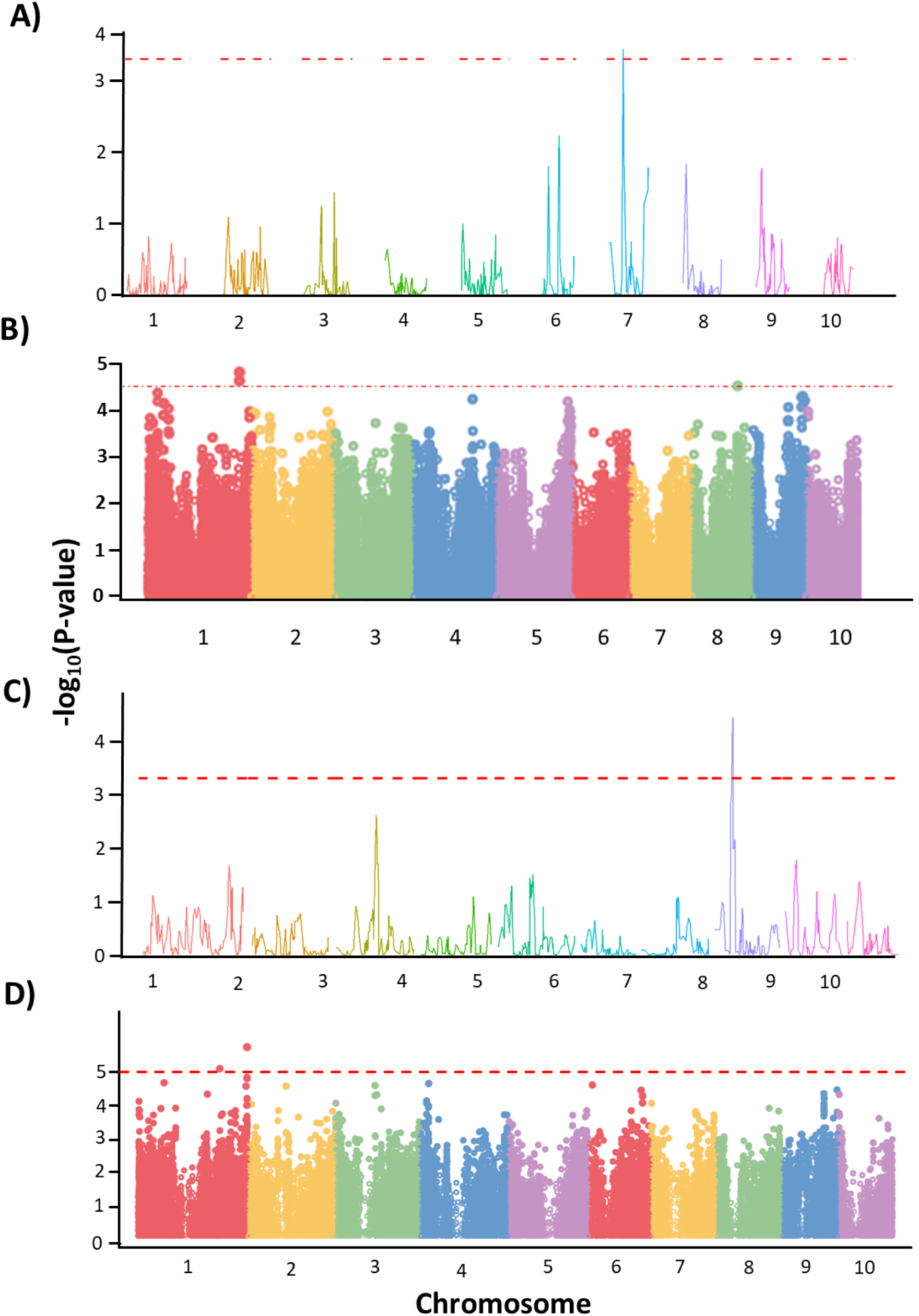
Genetic analysis for cortical cell length (CCL, A and B) and cortical cell diameter (CCD, C and D).QTL analysis results represented as test statistic −Log_10_(p) against marker genome location in IBM population for CCL (A) and CCD (C). GWAS analysis results represented as a Manhattan plot showing −Log_10_(p) value of markers in Widiv Panel for CCL (B) and CCD (D).

GWAS analysis using the BLINK method in GAPIT identified 23 significant SNPs (P < 0.00001) in nine genomic regions present on five different chromosomes (Figure 7B). The most significant SNPs lie on chromosome number 1 and were selected for further analysis. Candidate genes containing the significant SNPs were selected and the most significant SNP (rs1_270719512) was located in a gene model named Zm00001d028244. Zm00001d028244 encodes a hypothetical protein and is expressed throughout the plant.

QTL analysis on the IBM population reveals a significant QTL (named *QTL_CCD_8*) linked to CCD on chromosome number 8 (Figure 7C). SNP m851 and m853 demarcate (1.5 LOD interval) the *QTL_CCD_8* with m852 being the most significant SNP. Genotypes with homozygous BB allele of m852 have an average CCD of 45.63 µm, whereas homozygous AA allele genotypes have an average CCD of 39.97 µm, and heterozygous “AB” pool has the smallest diameter of 35.42 µm. The most significant marker (m852) in the *QTL_CCD_8* explains 24.23 % of the variation observed for CCD in the IBM mapping population.

GWAS analysis using the FarmCPU model identifies two significant SNPs (P < 0.00001) on chromosome 1 linked with CCD (Figure 7D). The most significant SNP is rs1_304962579 and the next significant SNP (rs1_304962580) is also in the LD block of rs1_304962579. The LD region (± 5000 Bp of rs1_304962579) around the significant SNP contains the Zm00001d034894 gene model which encodes ATP-dependent DNA helicase 2 subunit KU80.

### Phylogenetic analysis of CCD

Plant taxa were organized into five major groups including bryophytes – liverworts and mosses; lycophytes – the first vascular plants; and higher plants – gymnosperms and angiosperms. No clear phylogenetic trends in CCD were observed among these five groups (Figure 8). However, the average CCD in lycophyte, gymnosperm, and angiosperm groups was larger than bryophytes including liverwort and mosses but was not statistically significant (p = 0.86). It is also noteworthy that among these groups, no significant changes in minimum and maximum CCD were observed (Figure 8).

**Figure 8.**
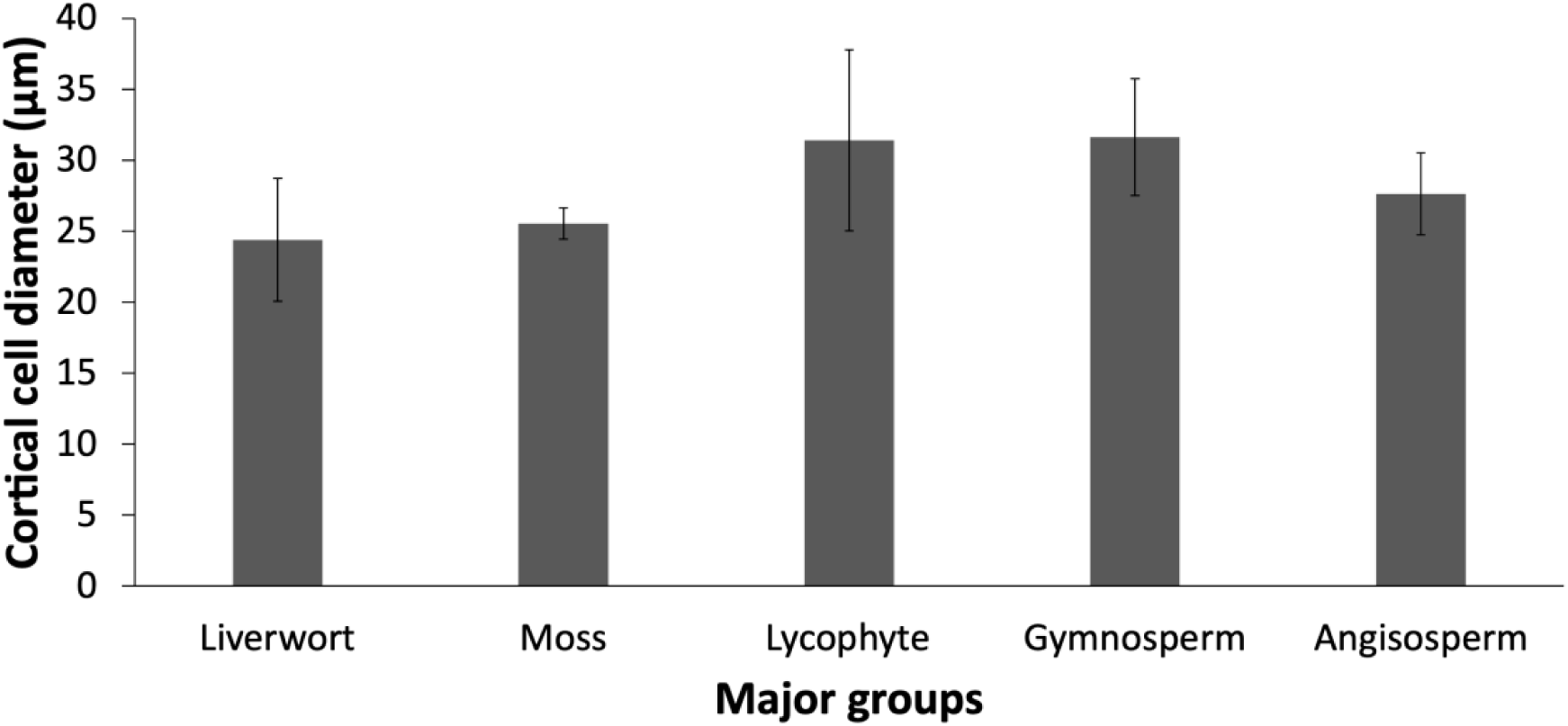
Variation for cortical cell diameter in the major evolutionary groups of plant species. Error bars indicate standard error centered on average cortical cell diameter. Number of samples in each group are as follows liverwort (7), moss (4), lycophyte (6), gymnosperm (15), and angiosperm (47). For details about the species in each group please refer to supplementary table S1.

## Discussion

In this study we describe how vacuolar occupancy and its relationship with cortical cell size (CCS) regulates root metabolic cost. Our results corroborate the hypothesis that an increased ratio of vacuolar to cytoplasmic volume is the underlying cause of reduced metabolic cost in root tissue with large CCS. We first show both cell diameter and cell length can determine CCS. By employing isophenic lines for CCD and CCL we show that an increase in either CCD or CCL is associated with reduced root respiration, nitrogen content, and phosphorus content. *RootSlice* modeling explicitly highlights the role of increased vacuolar occupancy in decreasing tissue nutrient content and mitochondrial density per unit cortical volume. The *RootSlice* predictions are validated by the SBFSEM imaging which shows that an increase in cell size either via an increase in cell diameter or cell length leads to an increase in vacuolar:cytoplasmic ratio per root volume. Utilizing GWAS and QTL analyses we identify QTLs and potential candidate genes underlying CCD and CCL which can be further explored for breeding purposes. Phylogenetic analysis reveals that no significant changes have occurred comparing CCD in different evolutionary groups including bryophytes, lycophytes, gymnosperms, and angiosperms. We propose CCD and CCL as potential phenes that can improve plant performance under conditions of suboptimal water and nutrient availability as well as soil mechanical impedance.

Root CCS is determined by cell diameter and cell length and both dimensions are functionally important for soil exploration. CCD defines cell cross-sectional area and contributes to root diameter (44). CCL determines the root elongation rate and its interplay with cell division determines root length (43). We observe significant variation for both CCD and CCL within maize germplasm (Figure 1A, B). As expected, the spread of variation for CCD is slightly larger within the maize diversity panel (∼1.6-fold) *WiDiv* (Figure 1A) as compared to the less diverse IBM-RIL population (∼1.5-fold, Figure 1A). Range and median CCD in the wheat Watkin collection are smaller than CCD in the maize populations (Figure 1). Smaller CCD in wheat is likely due to a smaller plant stature which allometrically relates to smaller diameter roots in wheat, while larger plant size in maize demands relatively thicker roots (52). However, the trends are the opposite for CCL, where we observe a 7.5-fold variation within the IBM-RILs (Figure 1B) but only a 3.35-fold variation within the *WiDiv* diversity panel (Figure 1B). Increased variation for CCL within the IBM-RIL population is mainly driven by the presence of RILs with reduced CCL (less than 40 µm), as the right tails of the distribution between *WiDiv* and IBM-RIL population are not significantly different. The extended left tail of the distribution for CCL within the IBM population may be related to inbreeding depression leading to shorter cells and less vigorous plants (53) Interestingly, the average CCL in the wheat Watkin collection is longer than both maize populations (IBM and WiDiv) (Figure 1B). This is likely due to different growing conditions where maize populations were grown in the field with greater soil mechanical impedance, but wheat CCL phenotyping was conducted using germination paper with negligible mechanical impedance to root growth.

It’s also noteworthy that variation for CCL is greater than variation for CCD in both the *WiDiv* diversity panel and IBM-RIL population. Increased range for CCL as compared to CCD is likely due to the role of cell length in root elongation rate and root length. Local soil taxa with constraints including mechanical impedance, water stress, and oxygen deficiency affect root growth (54). Therefore, these soil-specific properties may force plants to adapt specific root elongation rates and root length which are controlled partially by CCL, thereby leading to greater variation in CCL (54, 55). Similarly, comparing the diameter and length of maize lateral roots, Wu *et al*., 2016 show that the range of root length is significantly greater than the variation observed for root diameter (56). Cell length has also been shown to increase the size of vertebrate cardiac and skeletal muscle cells while the diameter does not change significantly (57).

Even though both phenes control cell size, no significant relationship between CCD and CCL was found within the WiDiv panel and wheat Watkin collection (Figure 1C) and a slight negative relationship was observed within the IBM-RIL population (Figure 1C). Therefore, CCL and CCD likely are independent phenes. The independence of CCD and CCL is also shown in our genetic analysis where no common QTLs were identified (Figure 7). CCD, unlike CCL, has been explored for its link with reduced root respiration. Chimungu et al., 2015 show that an increase in CCD is associated with reduced respiration and improved growth and yield under drought stress (23). Similarly, Colombi et al., 2019 show that wheat roots with large CCD have less energy demand and better soil penetration ability compared to roots with smaller CCD (24). Using our CCD isophenic lines and *RootSlice* modeling, we corroborate that increased CCD leads to reduced root respiration, nitrogen content, and phosphorus content. Therefore, increased CCD would potentially be beneficial in environments with suboptimal availability of nitrogen and phosphorus. By employing *OpenSimRoot* simulations Lopez-Valdivia *et al*., 2023 predicted the utility of larger cortical cells in maize under suboptimal nitrogen availability (27).

Unlike CCD, CCL has not been explored for its role in root metabolic cost. We show that increased CCL also reduces root respiration and tissue nutrient content. IBM-111, a genotype with longer cortical cells, has 53 % less respiration and 41 % less nitrogen content as compared to IBM-177, a genotype with shorter cortical cells. However, the magnitude of a decline in resource content with an increase in CCL is less than the magnitude of a decline in resource content with an increase in CCD. The difference in this magnitude is explained by cell geometry, with every unit increase in radius, r, cell volume increases by r^3^, whereas the relationship between cell volume and cell length is linear.

CCL can potentially improve plant performance under numerous edaphic stress conditions. CCL is likely associated with an increased root elongation rate (58) and increased metaxylem vessel length. Both increased root elongation rate and metaxylem vessel length can potentially improve drought tolerance by allowing efficient water acquisition from deeper soil domains. Therefore, increased CCL could be a novel phene for improving plant performance under suboptimal water and nutrient availability, which are primary constraints to plant growth in most terrestrial ecosystems (2, 6). Our CCL isophenic lines have utility for testing the utility of CCL under suboptimal water and nutrient conditions. Based on the variation observed within maize and wheat germplasm, it is probable that related crop species such as sorghum (*Sorghum bicolor*), millet (*Pennisetum glaucum*), barley (*Hordeum vulgare*), oat (*Avena sativa*), and rice (*Oryza sativa*) would also possess genotypic variation for CCL and CCD. Therefore, both CCL and CCD can be explored in any other monocot crop species for their benefits under edaphic stress.

Related to cortical parenchyma cells, size and number of parenchyma cells in the stele pith are other phenes that have not been explored in the context of root metabolic cost. We speculate that roots with larger and fewer pith parenchyma cells would have lower root metabolic cost and thus might be able to explore a larger volume of soil. Exploration of variation in pith parenchyma cells present in the diversity sets of major crops like maize and wheat would be a good start to understand the fitness landscape of stelar anatomy.

Examples of the negative relationship between cell size and metabolic cost, as described in this study, are rare in plants. However, several reports can be found in other biological taxa (31, 36). *E.g.* Schramm *et al.*, 2021 show that the Carabidae beetle species or different sexes with larger cells exhibit reduced mass-specific maintenance cost. Similarly, increased cell size in triploid *Xenopus* frogs reduces metabolic rate (59). In both cases, increased cell size reduces metabolic cost because of a reduction in total cell surface area which leads to reduced energetic cost of plasma membrane maintenance. However, this hypothesis does not completely explain the root cortical cell size-driven decrease in metabolic cost because along with the decrease in respiration, we also see a decrease in tissue nutrient content. The other possible explanation for the reduced root metabolic rate driven by the increased CCD and CCL is the increase in vacuolar: cytoplasmic ratio per root cortex volume, as previously proposed by Lynch, 2013 (45). To test this hypothesis, we employed *in silico* modelling using *RootSlice* and SBFSEM imaging.

*RootSlice* is a recent functional-structural model of root anatomy capable of accurately simulating root anatomical phenotypes (43). *RootSlice* simulates subcellular features including cell walls, plasma membrane, cytosol, vacuoles, and mitochondria. *RootSlice* is a heuristic model intended to explore hypotheses and the adequacy of conceptual models. Our employment of *RootSlice* in this study was aimed at testing the adequacy of the hypothesis that the empirically observed phenotypic variation in parenchyma cell size vacuolar volume could have significant influences on root metabolic costs. *RootSlice* permits manipulation of root cortical cell size by changing CCD and CCL independently. A previous study reports the use of *RootSlice* in showing the role of vacuole expansion in cell/root elongation (43) Therefore, *RootSlice* is a suitable platform to test the hypothesis of whether vacuolar occupancy in larger cells affects root metabolic cost.

For this study, *RootSlice* allowed us to simulate varying CCD and CCL phenotypes (Figure 4 and 5). The range of simulated CCD (10 to 25 µm) and CCL (30 to 300 µm) was defined to capture the natural variation observed for both phenes in maize. *In silico* results affirm that increased cell size, either by an increase in CCD or CCL, leads to a reduction in mitochondria density, respiration, tissue nitrogen, and phosphorus content (Figure 4 and 5). The magnitude of the decline in resource content with an increase in CCL is less than the magnitude of the decline in resource content with an increase in CCD. *RootSlice* results show that reduction in root metabolic cost is closely tracked by decreasing cytoplasmic volume per cortex volume (Figure 4C and 5C). The increase in vacuolar:cytoplasmic ratio per unit cortex volume accounts for decreased root respiration due to increased cell diameter and cell length (Figure 4 and 5). Reduced cytoplasmic volume leads to reduced mitochondrial density per unit cortex volume (Figure 4E and 5E). Reduction in phosphorus and nitrogen content is explained by reduced concentration of both nutrients in the vacuole compared to the symplasm (60, 61). In summary, results from *RootSlice* modeling are consistent with the hypothesis that vacuolar occupancy is sufficient to account for the empirically observed relationship of cortical cell size and root metabolic cost.

*RootSlice* modeling results were corroborated by SBFSEM imaging of cortical cells belonging to genotypes contrasting for CCD or CCL. SBFSEM enabled detailed imaging of the cytosol and vacuolar compartments while preserving the *in vivo* cell ultrastructure. However, the large diameter and long length of maize cells limited the imaging of whole cells and 3D-volume reconstruction. Thus, multiple images representing numerous cross-sectional and longitudinal sections of each cell were gathered. Surface area measurements of vacuolar and cytosolic compartments reveal a strong positive linear relationship between cell area and vacuolar area while the cytosolic compartment exhibits a weak relationship with cell size (Figure 6). Specifically, with each µm^2^ increase in cell surface area, vacuolar volume increased by 0.86 um^2^ while cytosolic area only increased by 0.14 um^2^. Therefore, increased cell size is mainly driven by an energy-efficient compartment *i.e.* vacuoles.

Genetic mapping employing GWAS and QTL analysis reveals multiple QTLs underlying CCD and CCL in maize (Figure 7). For CCL major QTL (QTL_CCL_7) was identified on chromosome 7 and for CCD major QTL (QTL_CCD_8) was identified on chromosome 8, however, low marker density in the IBM-RIL mapping population limited further annotation of the regions (62). GWAS analysis for both phenes resulted in higher resolution QTLs (Figure 7B and 7D). For CCL we identify numerous significant SNPs, the most significant of which is SNP rs1_270719512 on chromosome 1, which is associated with the gene model Zm00001d028244 encoding a hypothetical protein. An orthologue of this gene in rice (Os03g0221800) is annotated to be from the Leucine Rich Repeat (LLR) family. The LLR protein family is mainly known for its role in immunity (63) but some reports show a link with root morphogenesis and cell elongation (64). Furthermore, Zm00001d028244 is expressed throughout the plant with high expression in roots and meristematic zones. Expression in meristematic tissues supports the possibility of a role for this gene in cell morphogenesis, however, further confirmation is needed. For CCD, a significant SNP on chromosome 1 was associated with the Zm00001d034894 gene encoding an ATP-dependent DNA helicase 2 subunit KU80 protein. KU80 is well known for its role in DNA break repair but mutants for KU80 like proteins also exhibit abnormal cell size. Therefore, Zm00001d034894 could potentially be involved in regulating CCD. Confirmation of candidate genes and discovery of novel genes for CCD and CCL will help breeders to breed genotypes with increased CCD and CCL which might be useful under edaphic stress conditions.

To understand the evolutionary trajectory of root CCS, we measured CCD on numerous root cross-sectional images gathered from the literature. Measurements of CCL were limited by the unavailability of images of longitudinal sections of ancient plants. Plant species were grouped into major clades including rhizoid- bearing liverworts and mosses, vascular lycophytes, and higher plants grouped into gymnosperms and angiosperms. No clear pattern for CCD was observed among the different taxa spanning geologic time (Figure 8). We attribute this lack of variation to fitness tradeoffs accompanying the benefits of increased CCS. Tradeoffs of larger cells might include a higher cost of maintaining turgor pressure and reduced mechanical strength. The maintenance of turgor pressure in larger cells can be unfavorable in certain conditions such as osmotic/ salinity stress. For example Oertli (1986) showed that small epidermal cells were at least 20 times more resistant to negative turgor pressure-driven collapse as compared to large epidermal cells (65, 66). Another disadvantage of increased cell size could be reduced mechanical strength resulting from a larger ratio of symplasm volume to cell wall volume which can make larger cells more prone to rupturing or breaking under mechanical pressure or tension. Sapala et al., 2018 show that mechanical stress influences the fitness landscape of cell size and shape where smaller leaf epidermal cells are predicted to face less mechanical stress compared to larger cells (67).

Given the potential fitness benefits of increased CCS under drought stress, nutrient stress, and soil mechanical impedance, the benefit of decreased root metabolic cost might outweigh potential negative tradeoffs in modern agriculture. Global climate change is projected to intensify edaphic stress in the future (6), which may augment the utility of large CCS. It is striking that we observe large variation for CCD in maize germplasm but relatively low variability across plant phylogeny. This discrepancy in variation may be related to the small sample size for the phylogenetic analysis. We propose that a more detailed phylogenetic analysis of CCS is warranted.

In conclusion, we describe how vacuole occupancy in larger cortical cells can lead to a reduction in the metabolic cost of soil exploration. Increased CCD and CCL, both associated with vacuolar expansion, can reduce root maintenance costs and therefore may improve soil exploration and hence the acquisition of water and mineral nutrients. Although this study is primarily focused on maize and wheat, we propose that phenotypic variation in CCD and CCL would have comparable utility in other Poaceae species as well. Both CCD and CCL are under genetic control and thus we propose that both merit attention as ideotypes to breed crops resistant to edaphic stress.

## Materials and methods

### Germplasm selection

To select isophenic maize genotypes for CCD and CCL we utilized IBM (intermated B73 × M017) recombinant inbred lines (RILs). Being a RIL population descending from distinct F2 genotypes sharing the same genetic background, IBM is useful to select genotypes that can be potentially isophenic, reducing the risk of confounding effects (68). IBM has been phenotyped for many root anatomical and architectural phenes and a number of isophenic lines have been identified (62, 69). For this study, the IBM population (234 RILs) was grown under optimum field conditions at the Ukulima Root Biology Center (URBC) in Limpopo Province, South Africa (24°32.002’S, 28°07.427’E) from February through May 2011. At anthesis two representative root crowns from each plot were excavated and washed following the shovelomics pipeline. Two representative roots from the 4th whorl (for CCD) and 2^nd^ whorl (for CCL) of each crown were selected and 2 cm long roots were preserved in 75% ethanol in water (v/v) for anatomical analysis. Contrasting genotypes for CCD and CCL were identified including IBM 365 as larger CCD, IBM 30 as smaller CCD, IBM 111 as longer CCL, and IBM 177 as shorter CCL.

### Phenotyping CCD and CCL

To phenotype CCD and CCL, samples were sectioned and imaged using laser ablation tomography (LAT) as described by Strock *et al*., 2019 (70). For CCD, root cross-sections were imaged but for CCL, samples were ablated longitudinally. The cell diameter of each cortical cell file was measured to obtain CCD. The cell lengths of about 10 cells per sample were measured to obtain CCL. Unequal variance t-test was used to test determine whether average CCD and CCL was significantly different among isophenic sets of genotypes.

### Respiration and tissue nutrient content

Genotypes IBM 365, IBM 30, IBM 111, and IBM 177 were grown in greenhouse conditions. Each pot received 14 L of media which comprised of 50% (v/v) medium-grade sand (Quikrete Companies Inc., Harrisburg, PA, USA), 10% (v/v) soil (Ap2 Hagerstown silt loam (fine, mixed, semiactive, mesic Typic Hapludalf)), 20% (v/v) vermiculite (Whittemore Companies Inc., Lawrence, MA, USA), and 20% (v/v) perlite (Whittemore Companies Inc., Lawrence, MA, USA). Post-planting, each pot received 200 ml of nutrient solution daily, which consisted of (in µM): Ca(NO_3_)_2_ (1500), MgSO_4_ (1000), K_2_SO_4_ (500), Ca(H_2_PO_4_)_2_ (250), Fe-DTPA (Ferric sodium DTPA) (150), (NH_4_)_2_Fe(SO_4_)_2_ (100), H_3_BO_3_ (46.25), MnCl_2_ (9.14), ZnSO_4_ (0.76), CuSO_4_ (0.32), and (NH_4_)_6_Mo_7_O_2_ (0.08). Pots were arranged in a randomized complete block design with four blocks in total, each block including one replication of each treatment combination. Plants were grown for 40 days under a 16/8 h (light/dark) photoperiod, 40% relative humidity, and maximum/minimum temperatures of 28°C/26°C. Midday photosynthetic active radiation was approximately 900 to 1,000 μmol photons m^−2^ s^−1^. Natural light was supplemented from 06:00 to 22:00 with approximately 500 μmol photons m^−2^ s^−1^ from LED lamps. Upon harvesting, the root system was washed and root samples from the 4^th^ whorl (for CCD) and 2^nd^ whorl (CCL) were taken for root respiration, N content, and P content were taken. Root respiration rate measurements were conducted using a modified Li-Cor 6400 IRGA system following the protocol given by (71). For CCD, root N content was measured using an elemental analyzer (Perkin Elmer 2400 Series II CHN analyzer) and root P content was measured using ICP-MS (Perkin Elmer Avio 200 Optical Emission Spectrometer).

### *RootSlice* Simulations

For simulating varying CCD, we simulated root segments with 12 cortical cell files, a stele diameter of 356 µm, 0 % root cortical aerenchyma, CCL of 100 µm, 3 cell layers in the longitudinal direction, and 8 metaxylem vessels with an average metaxylem vessel diameter of 40 µm. By keeping the aforementioned parameters constant, we simulated four different CCD measurements including 10, 15, 20, and 25 µm.

For simulating varying CCL, we simulated root segments with 12 cortical cell files, a stele diameter of 356 µm, 0 % root cortical aerenchyma, CCD of 12 µm, 3 cell layers in the longitudinal direction, and 8 metaxylem vessels with an average metaxylem vessel diameter of 40 µm. By keeping the aforementioned parameters constant, we simulated 10 different CCL measurements including 30, 60, 90, 120, 150, 180, 210, 240, 270, and 300 µm.

### Serial block-facing scanning electron microscopy

All four maize genotypes (IBM 365, IBM 30, IBM 111, and IBM 177) were grown under in a growth chamber. Nutrient solution culture was used to obtain clean and intact root samples for serial block-facing scanning electron microscopy (SBFSEM). Each rep (4 reps in total) involving 4 plants (1 plant per genotype) was grown in a 20 L bucket filled with full-strength modified Barber’s nutrient solution which consisted of (in µM): Ca(NO_3_)_2_ (1500), MgSO_4_ (1000), K_2_SO_4_ (500), Ca(H_2_PO_4_)_2_ (250), Fe-DTPA (Ferric sodium DTPA) (150), (NH_4_)_2_Fe(SO_4_)_2_ (100), H_3_BO_3_ (46.25), MnCl_2_ (9.14), ZnSO_4_ (0.76), CuSO_4_ (0.32), and (NH_4_)_6_Mo_7_O_2_ (0.08) (72). Plants were grown for 40 d under a 16/8-h (light/dark) photoperiod, 40% relative humidity, and maximum/minimum temperatures of 28°C/26°C. Artificial light was provided from 06:00 to 22:00 with approximately 600 μmol photons m^−2^ s^−1^ using metal-halide lamps. Upon harvesting, representative roots from the 4th whorl (for CCD) and 2^nd^ whorl (for CCL) of each crown were selected and 0.5 cm long root segments were chemically fixed following the SBFSEM sample preparation protocol (73). Post-processing, samples were imaged using SBFSEM.

### Evolution of CCD

The goal for this part of the study was to evaluate CCD throughout terrestrial plant phylogeny. We assessed liverworts (*Marchantia polymorphera*), mosses (*Physcomitrella patens*), lycophytes (*Isoetes sp.*), gymnosperms (*Cycas taitugensis, Zamia integrifolia, Zamia vasquezii, Pinus pinaster, Pinus taeda, Welwitschia mirabilis*), and angiosperms (*Vaccinium corymbosum, Aspilia africana, Solanum lycopersicum, Salix sp., Elaeocarpus sp., Rumex crispus, Gunnera perpensa, Cucumis sativus, Glycine max, Betula platyphylla, Juglans microcarpa, Medicago tranculata, Pirus sp., Brassica juncea, Brassica napus, Phellodendron amurense, Acorus calamus, Maundia triglochinoides, Austrobaileya scandens, Cabomba caroliniana, Nuphar lutea, Amborella trichopoda, Elaeisguineensis sp., Eichhornia crassipes, Floscopa glabrata, Typhs glauca, Zingiber officinale, Xeronema callistemon, Saururus cernuus*). Further details about the specific species and literature used are given in the supplementary table S1. We gathered root cross-section images of different species representing each group. For liverworts and mosses, we measured the cell size in the rhizoids since these bryophytes lack true roots. For all the other groups, we were able to acquire true root images. CCD was measured for all the possible cortical cell files visible in an image and an average CCD measurement was attained.

### QTL analysis for CCD and CCL

QTL analysis for CCD and CCL was performed on the IBM RIL population. The IBM panel was grown at the Ukulima Root Biology Center in Alma, Limpopo, South Africa (24.32002°S, 28.07427°E) in 2011. The experimental design was a randomized complete block design with four replications. In each rep, each genotype was grown in a single-row plot consisting of 20 plants per plot. The distance between plants within a row was 23 cm and the row width was 75 cm. Optimal growing conditions were maintained with additional irrigation applied using a center pivot system as needed. A subset of 141 genotypes was used for this study. On these 141 genotypes, we have both phenotypic data for CCL and CCD as well as genotypic data for 1106 SNP markers.

Composite interval mapping was used to identify QTL with five marker covariates and a window size of 10 cM in R/qtl (74). The LOD threshold was determined using 1000 permutations at a significance threshold of 0.05. The Haley-Knott regression method was used to refine the positive effect of significant QTL. The physical position of markers is based on the version 4 B73 (AGPv4) reference sequence assembly (75). Further details regarding the IBM population and genetic maps are provided by Burton *et al.*, 2014 (76).

### GWAS for CCD and CCL

Genome-wide association study for CCD and CCL was performed using the *WiDiv* (Wisconsin diversity) panel. The panel was originally grown at the Apache Root Biology Center in Wilcox, AZ, USA (32.1539252°N, 109.4956928°W) in 2016 under optimal growing conditions (77). At anthesis, one representative plant per plot was excavated and a 3 cm segment from 5 to 8 cm from the basal portion of the root was excised from the crown for anatomical analysis. For this study, we re-phenotyped the fourth whorl root samples for CCD and CCL. The FarmCPU algorithm from the GAPIT R package was used to find marker-trait associations. Significant SNPs were identified based on a threshold of -log_10_(p-value) of 4. Significant SNPs identified in GWAS were translated to candidate genes based on the physical position of the genes in the version 4 B73 (AGPv4) reference sequence assembly (75). Expression analysis of the candidate genes was found in the MaizeGDB database (78).

### Wheat Watkin collection

The Watkin collection contains a diverse set of wheat landraces collected by A. E. Watkins in 1930 (79). For this study, we used a subset of 119 core landraces which represent most of the genetic diversity in the Watkin collection (80). To collect root anatomy samples, 10 seeds of each genotype were grown on a germination paper (Anchor Paper Co., St. Paul, MN, USA). After seed placement, each germination paper was rolled and kept in the germination chamber (set to 30 °C, no light, and 60-70 % humidity) for 48 hours. Upon germination (after 48 hours) the roll-ups were moved move to a growth chamber with a 16/8-h (light/dark) photoperiod (using artificial lighting), 40% relative humidity, and maximum/minimum temperatures of 24°C/26°C. After 7 days post-planting the plants were processed for root anatomy sample collection. From each roll-up (representing each genotype) four most healthy plants were selected and from each plant two most healthy seminal roots were taken for further processing. One cm away from the seed, 1.5 to 2 cm long root segments were harvested, placed into histocaps, and kept in 70 % EtoH. The samples were then dried using a critical point drier (Leica EM CPD 300, Leica Microsystems Inc, Buffalo Grove, IL, USA) to maintain the root structure. For imaging, we used laser ablation tomography (LAT) as described in detail by Strock et al., 2019 (70). CCD and CCL were measured using the ObjectJ macro in ImageJ software.

## Acknowledgments

We. acknowledge support from the US Department of Energy ARPA-E Award DE- AR0000821, the Howard G Buffett Foundation, and the US Department of Agriculture National Institute of Food and Agriculture and Hatch Appropriations Project PEN04732 and PENW-2020-03632.

## Data Availability statement

All the related data is available on Zenodo (DOI: 10.5281/zenodo.7709640).

