## Supplementary Data for "Cortical cell size regulates root metabolic cost"

This PDF file includes:

Supplementary Materials and Methods

Tables S1

Supplementary Table S1: Details about species included in the phylogenetic analysis of cortical cell diameter in the major evolutionary groups of plant species.

| ***Division*** | ***Sub-Division/ Major groups*** | ***Clade*** | ***Order*** | ***Species*** | ***Number of samples*** | ***References*** |
|---|---|---|---|---|---|---|
| *Bryophyte* | Liverwort | Liverwort | Marchantiales | *Marchantia polymorphera* | 7 | (1, 2) |
| *Bryophyte* | Moss | Moss | Funariales | *Physcomitrella patens* | 4 | (3) |
| *Lycophyte* | Lycophyte | Lycophyte | Isoetales | *Isoetes sp.* | 6 | (4) |
| *Spermophyte* | Angisosperm | Eudicot | Ericales | *Vaccinium corymbosum* | 2 | (5) |
| *Spermophyte* | Angisosperm | Eudicot | Asterales | *Aspilia africana* | 1 | (6) |
| *Spermophyte* | Angisosperm | Eudicot | Solanales | *Solanum lycopersicum* | 2 | (7) |
| *Spermophyte* | Angisosperm | Eudicot | Malpighiales | *Salix sp.* | 2 | (8) |
| *Spermophyte* | Angisosperm | Eudicot | Oxalidales | *Elaeocarpus sp.* | 4 | (9) |
| *Spermophyte* | Angisosperm | Eudicot | Cryophyllales | *Rumex crispus* | 1 | (10) |
| *Spermophyte* | Angisosperm | Eudicot | Gunnerales | *Gunnera perpensa* | 2 | (11) |
| *Spermophyte* | Angisosperm | Eudicot | cucurbitales | *Cucumis sativus* | 1 | (12) |
| *Spermophyte* | Angisosperm | Eudicot | cucurbitales | *Glycine max* | 1 | (13) |
| *Spermophyte* | Angisosperm | Eudicot | Fagales | *Betula platyphylla* | 1 | (14) |
| *Spermophyte* | Angisosperm | Eudicot | Fagales | *Juglans microcarpa* | 1 | (15) |

| | | | | | | |
|---|---|---|---|---|---|---|
| *Spermophyte* | Angisosperm | Eudicot | Fagales | *Medicago tranculata* | 3 | (16) |
| *Spermophyte* | Angisosperm | Eudicot | Rosales | *Pirus sp.* | 1 | (8) |
| *Spermophyte* | Angisosperm | Eudicot | Brassicales | *Brassica juncea* | 1 | (17) |
| *Spermophyte* | Angisosperm | Eudicot | Brassicales | *Brassica napus* | 1 | (17) |
| *Spermophyte* | Angisosperm | Eudicot | Sapindales | *Phellodendron amurense* | 1 | (14) |
| *Spermophyte* | Angisosperm | Monocots | Acorales | *Acorus calamus* | 2 | (18) |
| *Spermophyte* | Angisosperm | Monocots | Alismatales | *Maundia triglochinoides* | 1 | (19) |
| *Spermophyte* | Angisosperm | Monocots | Austrobaileyales | *Austrobaileya scandens* | 2 | (20) |
| *Spermophyte* | Angisosperm | Monocots | Nymphaeales | *Cabomba caroliniana* | 1 | (10) |
| *Spermophyte* | Angisosperm | Monocots | Nymphaeales | *Nuphar lutea* | 1 | (10) |
| *Spermophyte* | Angisosperm | Monocots | Amborellales | *Amborella trichopoda* | 4 | (10) |
| *Spermophyte* | Angisosperm | Monocots | Arecales | *Elaeisguineensis sp.* | 4 | (21) |
| *Spermophyte* | Angisosperm | Monocots | commelinales | *Eichhornia crassipes* | 1 | (10) |
| *Spermophyte* | Angisosperm | Monocots | commelinales | *Floscopa glabrata* | 2 | (22) |
| *Spermophyte* | Angisosperm | Monocots | poales | *Typhs glauca* | 1 | (10) |
| *Spermophyte* | Angisosperm | Monocots | zingibirales | *Zingiber officinale* | 1 | (23) |
| *Spermophyte* | Angisosperm | Monocots | Asparagales | *Xeronema callistemon* | 1 | (24) |
| *Spermophyte* | Angisosperm | Monocots | Piperales | *Saururus cernuus* | 1 | (10) |
| *Spermophyte* | Gymnosperm | Cycadales | Cycadales | *Cycas taitugensis* | 2 | (25) |
| *Spermophyte* | Gymnosperm | Cycadales | Cycadales | *Dioon edule* | 1 | Gutiérrez-Arroyo *et al*., unpublished |
| *Spermophyte* | Gymnosperm | Cycadales | Cycadales | *Zamia integrifolia* | 1 | (25) |
| *Spermophyte* | Gymnosperm | Cycadales | Cycadales | *Zamia vasquezii* | 1 | (25) |
| *Spermophyte* | Gymnosperm | Pinales | Pinales | *Pinus pinaster* | 2 | (26) |
| *Spermophyte* | Gymnosperm | Pinales | Pinales | *Pinus taeda* | 7 | (27) |
| *Spermophyte* | Gymnosperm | Welwitschiales | Welwitschiales | *Welwitschia mirabilis* | 1 | (28) |
